# Chemically informed representations of amino acids enable learning beyond the canonical protein alphabet

**DOI:** 10.64898/2026.03.12.711352

**Authors:** Jialu Chen Christiansen, Miguel González-Valdés Tejero, Chuang Sun Hembo, Yuchen Li, Carolina Barra

## Abstract

Computational models of proteins typically represent sequences using a fixed twenty-letter alphabet describing canonical amino acids. Although this symbolic representation underlies most machine learning approaches to protein analysis, it abstracts away the chemical structure of residues and cannot naturally encode post-translational modifications (PTMs). As a result, current models struggle to incorporate chemical variation beyond the canonical amino acid alphabet.

Here we introduce a chemically informed representation of peptides based on two-dimensional depictions of amino acid structures. Peptides are encoded as mosaics of residue depictions and embedded using a convolutional autoencoder, allowing machine learning models to learn physicochemical features directly from molecular structure. Because these representations capture chemical properties rather than symbolic residue identities, they enable learning across structurally related residues and support generalization to modified amino acids not explicitly observed during training. Applied to Major Histocompatibility Complex class I binding prediction, these embeddings achieve competitive performance while enabling chemically interpretable attribution of the molecular features driving predictions.

## Introduction

The language of proteins is commonly represented using a twenty-letter amino acid alphabet, a simplification that has enabled major advances in bioinformatics but abstracts away the underlying chemical structure of proteins. This symbolic representation allowed proteins to be conceptualized as strings, facilitating the development of foundational bioinformatics tools, ranging from sequence alignment algorithms (1,2,3) and substitution matrices such as BLOSUM and PAM (4,5), to large-scale comparative genomics and modern protein structure prediction methods (6). More recently, machine learning approaches have built upon these sequence-based encodings, evolving from simple one-hot representations (7) and physicochemical feature vectors (8,9) to deep neural networks and large protein language models (10,11) capable of capturing evolutionary and structural relationships across millions of sequences. Together, these advances have transformed our ability to analyze protein function and interactions directly from sequence information.

Despite these advances, most computational models still rely on symbolic sequence representations that cannot explicitly capture the chemical diversity of amino acids and their post-translational modifications. As a result, current machine learning frameworks struggle to incorporate chemical variation beyond the canonical amino acid alphabet, limiting their ability to model protein behaviour in realistic biological contexts.

Protein function is strongly influenced by physicochemical context and chemical modification. Environmental factors such as pH and ionic strength alter protonation states, electrostatic interactions, and protein stability (12,13), while post-translational modifications (PTMs) including phosphorylation, glycosylation, ubiquitination, acetylation, and citrullination substantially expand the functional diversity of the proteome (14–16). These covalent modifications alter the charge, steric properties, conformational dynamics, and binding affinity of proteins, thereby regulating processes such as signal transduction, transcription, metabolism, immune recognition, and protein degradation (17). For example, phosphorylation can rapidly switch protein activity on or off and is central to many signaling pathways (18); ubiquitination commonly targets proteins for degradation via the proteasome and thus controls protein lifetime and cell-cycle progression (19); glycosylation modulates protein folding, stability, and cell-cell interactions important in immune recognition and disease (20); and lysine acetylation regulates transcriptional activity, chromatin structure, and metabolic responses (21). Because these modifications ultimately arise from changes in molecular structure, representing amino acids through chemically explicit depictions rather than symbolic letters may allow machine learning models to learn transferable physicochemical principles and to generalize to chemically modified residues that were not explicitly present during training.

This limitation persists even in modern protein language models that learn thousands or millions of contextual features from large evolutionary sequence datasets (10,22–23). Although these models capture rich evolutionary patterns, residues are still encoded exclusively as canonical amino acids. Consequently, chemically modified residues cannot be represented directly. When an amino acid undergoes modification—for example, phosphorylation, which affects roughly one third of human proteins (24)—these models cannot capture the resulting changes in charge, steric properties, or interaction potential. Several attempts have been made to address this limitation. In peptide-MHC binding prediction, for example, the amino acid alphabet has been extended to include additional symbols representing phosphorylated serine (B), threonine (Z), or tyrosine (O) (25). Although phosphorylated serine can mimic the charge and steric properties of glutamic acid and function as an alternative binding anchor, such extensions remain ad hoc and do not generalize across the wide diversity of PTMs found in biological systems (26). As a result, modified residues are often ignored, replaced, or simplified when sequences are used as input for machine learning models, leaving a significant gap in computational approaches capable of representing the chemical diversity of proteins.

The ability to represent modified residues is particularly important in immune recognition. Chemically modified self-proteins can generate neo-epitopes that escape immune tolerance and trigger pathological immune responses in autoimmune disease. Increasing evidence shows that several autoimmune disorders are driven by immune recognition of modified self-antigens rather than their canonical counterparts (27–29). For example, the enzymatic conversion of arginine to citrulline alters protein charge and structure, generating epitopes that stimulate the production of anti-citrullinated protein antibodies, a hallmark of rheumatoid arthritis (30). More broadly, post-translationally modified peptides presented by major histocompatibility complex (MHC) molecules can shape autoreactive T-cell responses in diseases such as type 1 diabetes and inflammatory rheumatic disorders (31,32). Recent studies combining immunopeptidomics and structural immunology further demonstrate that modified peptides can alter antigen presentation and T-cell receptor recognition, expanding the repertoire of potential autoantigens (31,33). These findings highlight the need for computational models capable of representing chemically modified residues when studying protein-peptide interactions and immune recognition.

Recent advances in deep learning and computer vision provide an opportunity to address this challenge. Convolutional architectures such as AlexNet, VGG, ResNet, and Inception, together with generative models including variational autoencoders, have demonstrated powerful capabilities for extracting hierarchical features from complex visual inputs (34–38). At the same time, cheminformatics toolkits can now generate standardized two- and three-dimensional depictions of molecular structures from chemical descriptions such as SMILES strings (39,40). These developments make it possible to represent amino acids not as abstract symbols but as explicit chemical structures that encode their physicochemical properties.

Here we introduce a chemically informed representation paradigm that replaces symbolic amino acid letters with explicit structural depictions of residues. Individual amino acids are represented through standardized two-dimensional molecular images that inherently encode physicochemical properties such as charge distribution, steric size, hydrophobicity, and functional groups. Because canonical and chemically modified residues share underlying structural features, these representations allow machine learning models to learn physicochemical relationships between residues rather than relying solely on symbolic identity. This framework provides a basis for modeling post-translational diversity and enables models to infer the behaviour of chemically modified residues that were not present during training. In addition, representing residues as molecular depictions enables the use of visual attribution methods that map predictive signals directly onto molecular structures, providing an interpretable way to assess which chemical features drive model predictions.

To evaluate this framework, we generated standardized two-dimensional depictions of amino acid structures and assembled them into peptide mosaics that preserve both structural and sequential information. A convolutional autoencoder was used to learn compact latent embeddings from these peptide depictions, capturing structural features of residues and their spatial arrangement. We then evaluated these representations in the task of predicting peptide binding to MHC class I molecules, a central problem in immunology and vaccine design, and compared their performance with conventional one-hot encodings and sequence-similarity baselines using nested cross-validation across multiple Human Leukocyte Antigen (HLA) alleles. Our results showed that image-derived peptide embeddings supported competitive classification performance, particularly in the low false-positive regime, while enabling chemically interpretable attribution of the molecular features driving peptide-MHC interaction predictions.

## Methods

### Dataset construction and preprocessing

Peptide binding data were derived from publicly available immunopeptidomics datasets containing experimentally identified MHC class I ligands including phosphorylations. The dataset was filtered to retain only human peptides of length nine residues, producing an initial dataset denoted D_Human9, where each entry consisted of a peptide sequence, a binding label, and the originating cell line. Non-human peptides and sequences with ambiguous annotations were removed during preprocessing to ensure a consistent evaluation set.

To assign peptides to specific HLA alleles, peptides were reannotated using the MHCMotifDecon tool (version 1.0). This tool deconvolutes peptide motifs derived from cell-line datasets and assigns peptides to the most likely presenting HLA allele. Prior to submission, phosphorylated residues were replaced with the placeholder symbol “X”, as the tool does not accept non-standard amino acids. Allele information was provided through the reference allele lists distributed with the original dataset. The resulting dataset, denoted D_HLA9, contained entries formatted as peptide sequence, binding label, and assigned HLA allele.

To incorporate structural context from the MHC binding groove, an extended dataset was generated by concatenating peptide sequences with corresponding MHC pseudo-sequences. Pseudo-sequences were obtained from the NetMHCpan-4.1 dataset archive and represent key polymorphic residues within the peptide-binding region of each HLA molecule. By aligning cell-line identifiers with the pseudo-sequence reference file, a combined dataset D_HLA9P was generated in which each input consisted of the peptide sequence followed by its associated MHC pseudo-sequence together with the binding label.

The datasets D_Human9, D_HLA9, and D_HLA9P were used consistently throughout the study for training and evaluation of peptide representations and prediction models.

### Amino acid structural representations

To generate chemically informed representations of amino acids, we constructed standardized two-dimensional depictions of molecular structures using RDKit. Canonical isomeric Simplified Molecular Input Line Entry System (SMILES) strings for the twenty standard amino acids, as well as phosphorylated variants of serine, threonine, and tyrosine, were retrieved from PubChem and converted into RDKit molecule objects. Two-dimensional coordinates were then computed for each molecule and aligned to a common peptide backbone template. This alignment ensured a consistent orientation of side chains across residues and reduced rotational variance in the generated images. The resulting depictions provide a uniform structural representation of amino acids while preserving information about atomic connectivity, functional groups, and overall molecular geometry.

### Peptide mosaic construction

Whole peptides were represented by concatenating the individual amino acid depictions along the horizontal axis in sequence order. Each residue was rendered as a small image representing its chemical structure, and the ordered concatenation of these images produced a two-dimensional mosaic encoding both structural and sequential information. In this representation, each residue retains its molecular depiction while the overall arrangement reflects the primary sequence of the peptide. This approach produces array-like image representations of peptides that simultaneously preserve chemical features of residues and positional relationships within the sequence.

### Learning peptide embeddings

Peptide mosaics were processed using a convolutional autoencoder designed to learn compact feature representations from the structural images. The encoder consisted of four convolutional blocks, each including a 3×3 convolution with padding, batch normalization, a LeakyReLU activation function (slope 0.1), and 2×2 max pooling. These operations progressively reduced the spatial dimensions of the peptide image and produced a compressed feature map. The resulting representation was flattened and passed through a fully connected layer to generate a 256-dimensional latent vector representing the peptide. A mirrored decoder composed of transpose convolution and upsampling layers reconstructed the input image from the latent representation. The autoencoder was trained using mean squared error loss between the input and reconstructed peptide images. In this framework, the encoder functions as a trainable feature extractor that learns latent representations of peptide structure.

### Baseline sequence encodings

To provide a baseline for comparison with the image-based representation, peptide sequences were also encoded using conventional one-hot encoding. In this representation, each amino acid was mapped to a binary vector of length 23, corresponding to the twenty canonical amino acids plus three phosphorylated residues. Phosphorylated serine, threonine, and tyrosine were represented as distinct tokens to distinguish them from their unmodified counterparts. A peptide sequence of length nine was therefore encoded as a 9×23 binary matrix, which was subsequently flattened into a feature vector prior to classification. This encoding treats residues as discrete categories and provides a standard reference representation widely used in peptide-MHC binding prediction tasks.

### MHC class I binding prediction task

The learned representations were evaluated using the task of predicting peptide binding to MHC class I molecules. The dataset used here consisted only of nine-mer peptides annotated with binding labels and associated HLA alleles. Peptides derived from cell-line datasets were reassigned to their corresponding HLA alleles using the MHCMotifDecon tool, and the resulting dataset was used to construct peptide-allele pairs for training and evaluation.

### Model training and evaluation

The overall modeling pipeline consisted of two stages. First, the convolutional autoencoder was trained in an unsupervised manner to reconstruct peptide depictions and learn latent embeddings. The trained encoder component was then frozen and used to extract fixed-length feature vectors for each peptide. In the second stage, these embeddings were used as input to a feedforward neural network classifier for MHC class I binding prediction. Model development and evaluation were performed using nested cross-validation, with five outer folds for performance estimation and inner training/validation splits for model selection. The outer test fold was excluded entirely from representation learning, classifier training and validation at each iteration.

### Attribution and interpretability analysis

To investigate which molecular features contributed to model predictions, attribution analyses were performed on the image-based peptide representations. Gradient-based saliency methods were used to highlight regions of the input image that most strongly influenced model predictions. In the context of molecular depictions, these attribution maps can be projected directly onto the chemical structures of residues, enabling identification of substructures or functional groups that contribute to predicted binding. These analyses were used qualitatively to assess whether learned representations reflected chemically meaningful features rather than artifacts of the visual representation.

## Results

To evaluate whether chemically informed peptide representations can support machine learning models of peptide-MHC interactions, we first constructed image-based depictions of amino acids and assembled them into peptide mosaics. We then trained a convolutional autoencoder to learn compact latent embeddings from these structural representations. The resulting embeddings were evaluated in the task of predicting peptide binding to MHC class I molecules and compared with conventional sequence encodings. Finally, we analyzed model attribution patterns and dataset characteristics to assess the interpretability and limitations of the learned representations, including their ability to represent chemically modified residues. Because these representations encode the chemical structure of amino acids rather than symbolic residue identities, they provide a potential framework for modeling chemically modified residues within the same representation space.

### Image-based representations capture structural features of amino acids

To construct chemically informed peptide representations, individual amino acids were rendered as standardized two-dimensional depictions of their molecular structures and concatenated to form peptide mosaics. These depictions preserve key structural characteristics of residues, including side-chain size, charged or charge-associated functional groups, and bond connectivity. Alignment of residues to a common peptide backbone ensured consistent orientation across images, enabling the model to compare structural features across different amino acids. Representative examples of amino acid depictions are shown in Fig. 1A, illustrating how side-chain structures are preserved within the standardized representation. Whole peptides were then constructed by concatenating these residue depictions along the sequence axis to produce mosaic-like peptide images (Fig. 1B). Visual inspection confirmed that the resulting peptide mosaics retained both the sequential arrangement of residues and the structural features of individual amino acids. Modified residues, such as phosphorylated serine, threonine, and tyrosine, displayed clear structural differences from their canonical counterparts, demonstrating that the representation can explicitly encode post-translational modifications.

**Figure 1.**
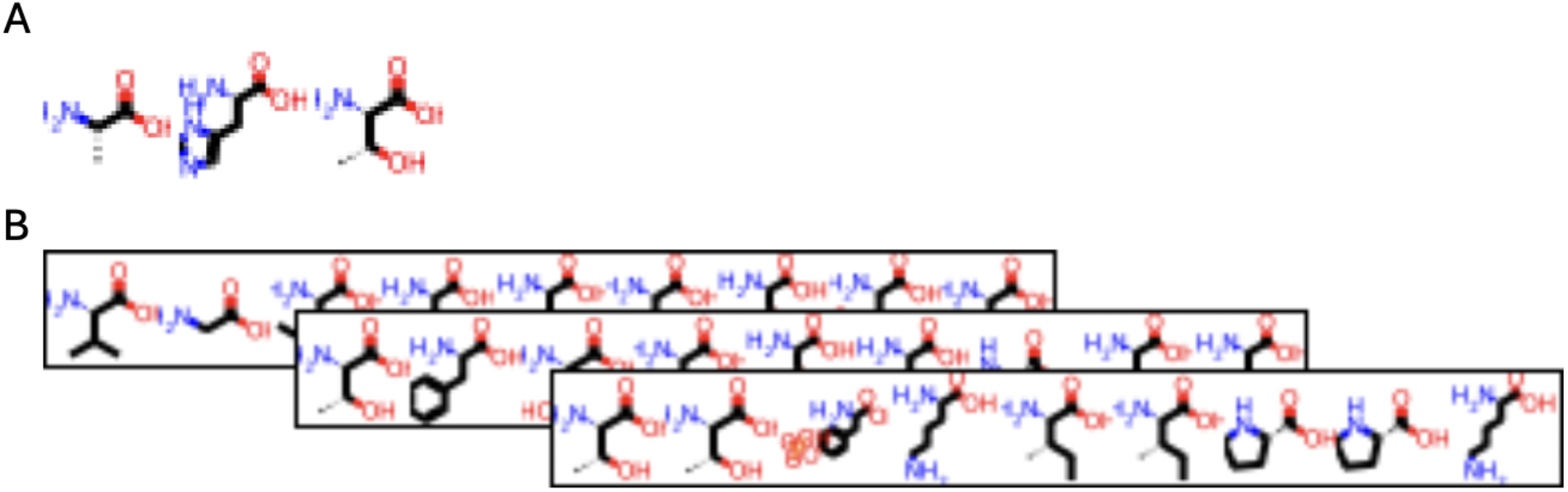
Chemically informed peptide representations based on amino acid structural depictions. **A**, Standardized two-dimensional depictions of representative amino acids generated from SMILES strings using RDKit. Each residue is aligned to a common peptide backbone template to ensure consistent orientation across side chains while preserving molecular features such as bond connectivity, functional groups, and side-chain geometry. **B**, Example peptide mosaics constructed by concatenating individual amino acid depictions along the sequence axis. This representation preserves the sequential order of residues while encoding their underlying chemical structure, allowing peptides to be represented as image-like inputs suitable for convolutional neural networks. Modified residues, such as phosphorylated amino acids, can be incorporated directly within this framework through their corresponding molecular depictions.

Because these representations encode the chemical structure of amino acids directly, they provide a representation space in which chemically modified residues can be incorporated without expanding the symbolic amino acid alphabet.

### Autoencoder models learn compact peptide embeddings

To learn compact representations of the peptide mosaics, a convolutional autoencoder was trained to reconstruct peptide depictions from compressed latent vectors. The architecture of the autoencoder is illustrated in Fig. 2A. The encoder progressively reduced the spatial dimensions of the peptide images through convolutional and pooling layers before projecting the features into a 256-dimensional latent space. Reconstruction results indicated that the model was able to capture the overall structural arrangement of peptide depictions. Representative examples of reconstructed peptide mosaics are shown in Fig. 2B,C, where the reconstructed images closely resemble the original inputs when trained on sufficiently large datasets. These results suggest that the learned latent representations preserve key structural characteristics of the peptides. However, reconstruction quality decreased when the model was trained on smaller datasets, indicating that the stability of the learned representation depends on data availability.

**Figure 2.**
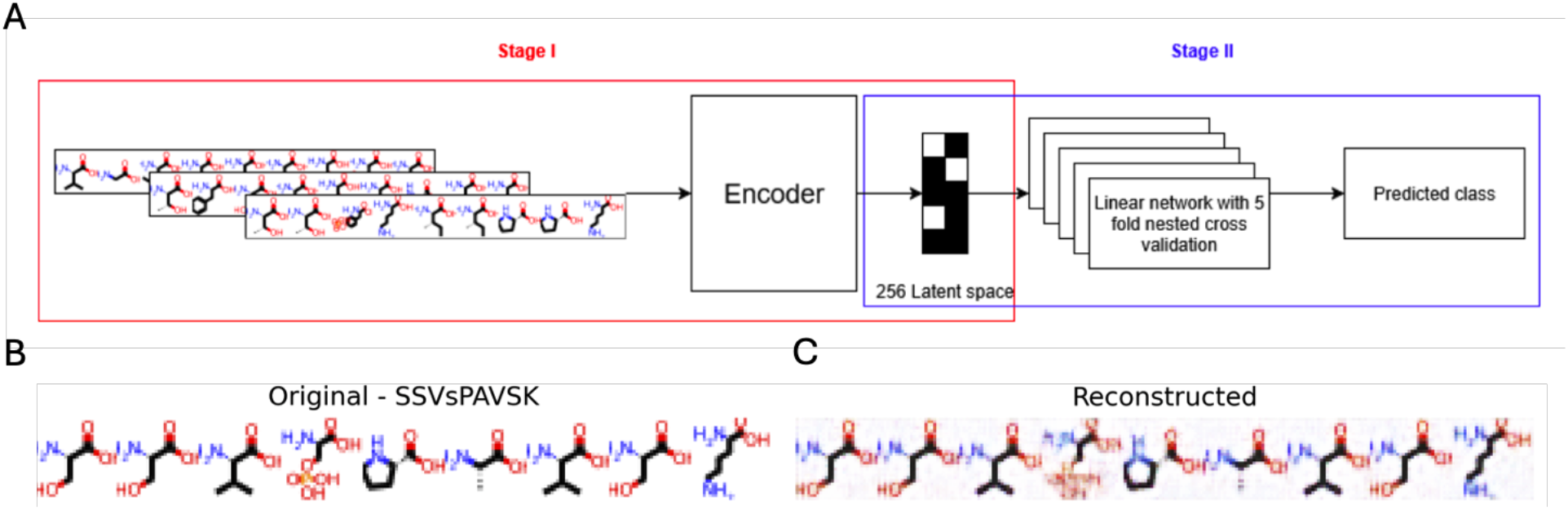
Convolutional autoencoder for learning peptide representations from structural depictions. **A**, Overview of the two-stage modeling framework used in this study. In Stage I, peptide mosaics generated from concatenated amino acid depictions are processed by a convolutional autoencoder encoder that compresses the input representation into a 256-dimensional latent vector capturing structural features of the peptide. In Stage II, these latent embeddings are used as input features for a linear classifier trained with nested cross-validation to predict peptide binding to Major Histocompatibility Complex class I (MHC-I) molecules. **B**, Example input peptide mosaics used to train the autoencoder. The lower case s represents the phosphorylated serine. **C**, Corresponding reconstructed peptide mosaics generated by the decoder. The reconstructed images preserve the overall structure and arrangement of amino acid side chains, indicating that the latent representation captures key structural characteristics of the peptide inputs.

Because these representations encode the underlying chemical structure of amino acids, they provide a foundation for modeling chemically modified residues within the same representation space.

### Image-derived embeddings support peptide-MHC binding prediction

The learned latent representations were evaluated in the task of predicting peptide binding to MHC class I molecules (Fig. 2A, Stage II). A feedforward neural network classifier was trained using the autoencoder-derived peptide embeddings as input features. Predictive performance was assessed across multiple HLA alleles using the area under the receiver operating characteristic curve (AUC).

The resulting performance across alleles is summarized in Fig. 3, which compares the AUC achieved by models trained on image-derived peptide embeddings with those trained on conventional one-hot encoded sequences. The autoencoder-based representation achieved meaningful predictive performance across most alleles and consistently outperformed simple sequence similarity approaches such as BLAST (not shown). However, models trained on one-hot encoded sequences generally achieved higher AUC values, reflecting the strong contribution of position-specific residue identities in peptide-MHC binding prediction.

**Figure 3.**
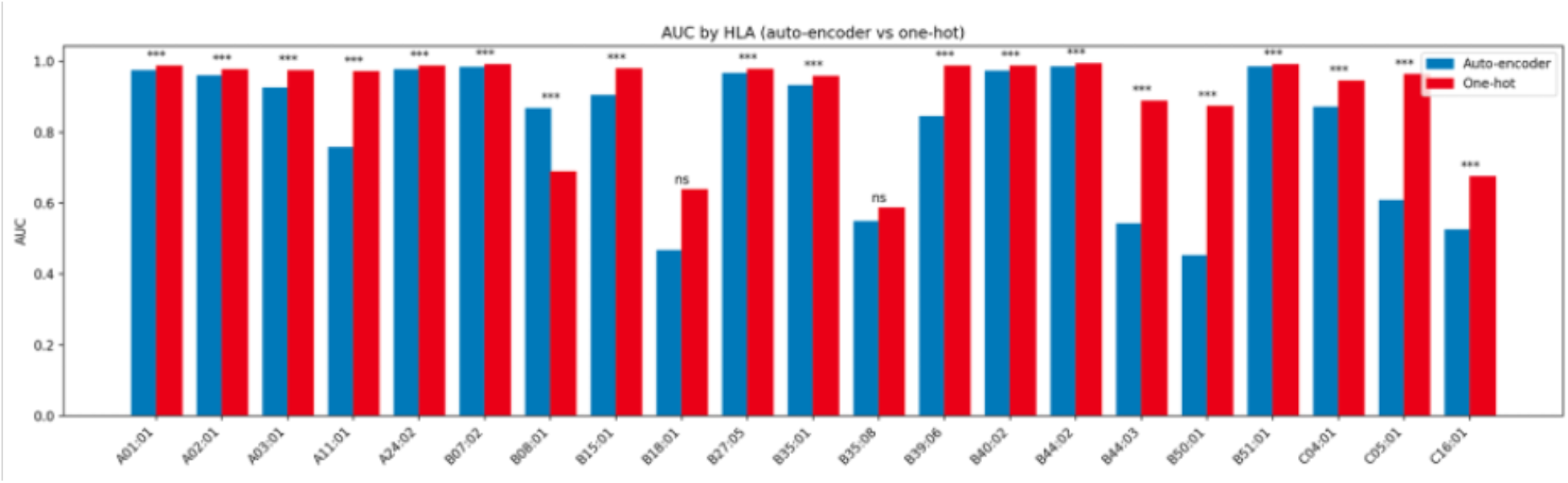
Prediction performance across HLA alleles using image-derived peptide embeddings. Area under the receiver operating characteristic curve (AUC) for peptide-MHC class I binding prediction across individual HLA alleles. Bars show the predictive performance of classifiers trained on autoencoder-derived peptide embeddings (blue) compared with models trained on conventional one-hot encoded peptide sequences (red). Each pair of bars corresponds to a specific HLA allele. Statistical significance of performance differences between encoding strategies is indicated above each pair of bars (***P < 0.001; ns, not significant). While one-hot encodings generally achieve higher AUC values, the image-derived representations maintain competitive predictive performance across several alleles, indicating that structural peptide representations capture features relevant to peptide-MHC binding.

Despite this difference, the image-derived embeddings achieved competitive predictive performance across several alleles and retained meaningful signal throughout the dataset. These results demonstrate that structurally informed peptide representations capture biologically relevant features of peptide–MHC interactions while learning patterns that extend beyond explicit residue identity. Importantly, because the proposed encoding represents amino acids through their underlying chemical structure rather than symbolic labels, the learned representations provide a framework for modeling chemically modified residues without requiring additional tokens in the sequence alphabet.

### Generalization to unseen post-translationally modified peptides

To test whether the learned representations capture the underlying chemical properties of amino acids and can therefore generalize to unseen modifications, we evaluated the model on peptides containing phosphorylated residues that were not explicitly observed during classifier training. Peptide mosaics were generated in which phosphorylated serine, threonine, and tyrosine were represented using their corresponding molecular depictions. Latent embeddings were then extracted using the trained encoder and used as input for the downstream feedforward neural network classifier.

We focused on the allele HLA-B*40, whose peptide-binding motif is characterized by two primary anchor positions: glutamic acid or aspartic acid at position 2 (P2) and methionine, phenylalanine, or other aliphatic residues at the C-terminus. Previous studies have shown that phosphorylation of serine at P2 can mimic the negatively charged side chains of glutamic acid or aspartic acid and may therefore substitute for these residues at the anchor position. Indeed, several phosphopeptides harboring pSer at P2 have been experimentally identified as ligands of HLA-B*40 (41).

We therefore evaluated whether peptides containing phosphorylated serine at P2, an amino acid modification not previously encountered by the classifier,could be correctly predicted as binders of HLA-B40. Notably, these peptides were predicted as binders for HLA-B40 but not for alleles with different anchor residue preferences at P2. These observations suggest that the chemically informed representation enables the model to recognize structural and physicochemical similarities between phosphorylated residues and canonical anchor residues, allowing generalization to modified amino acids not explicitly present during training.

### Attribution analysis reveals chemically interpretable features

To further investigate which structural features contributed to model predictions, attribution analyses were performed on the image-based peptide representations. Gradient-based saliency methods were applied to the trained classifier to identify regions of the input peptide depiction that most strongly influenced the predicted binding outcome. An example of this analysis is shown in Fig. 4, where the original peptide mosaic contains a phosphorylation at position 2 (Fig. 4A) is compared with the attribution overlay (Fig. 4B) and the corresponding attribution heatmap (Fig. 4C). These attribution maps highlight localized regions of the peptide depiction that contribute most strongly to the prediction, enabling interpretation of the model’s decision in terms of specific structural elements of the peptide.

**Figure 4.**
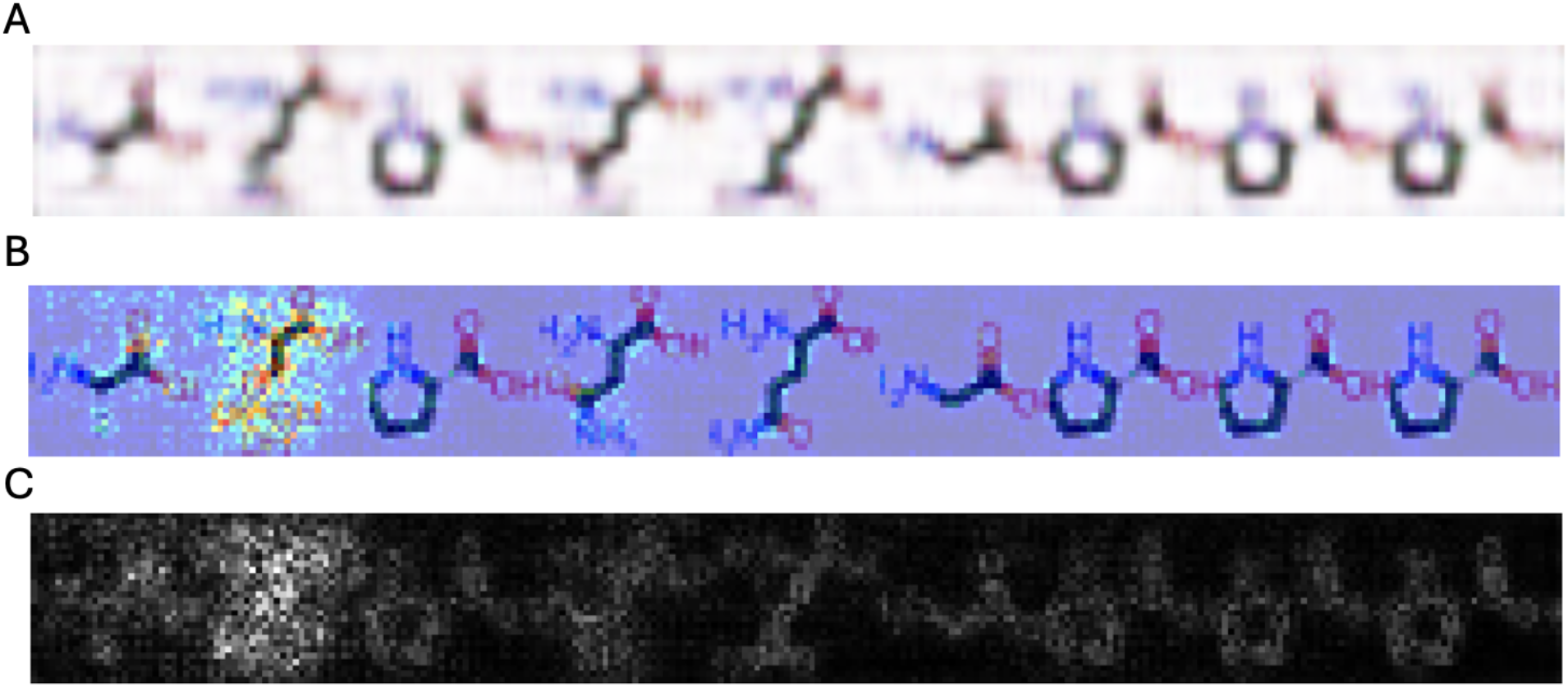
Attribution analysis reveals regions of peptide depictions contributing to model predictions. **A**, Example decoder reconstruction of an unseen phosphorylated peptide. The mosaic is constructed by concatenating two-dimensional depictions of individual amino acids along the sequence axis. **B**, Overlay of the attribution map on the peptide mosaic, highlighting regions of the input image that most strongly influence the model prediction. Warmer colors indicate higher attribution scores. **C**, Corresponding attribution heatmap shown independently of the peptide depiction. Brighter regions indicate image locations with higher influence on the prediction. The attribution maps highlight localized regions corresponding to specific amino acid side chains and positions within the peptide, demonstrating how the image-based representation enables visualization of structural features that contribute to peptide-MHC binding predictions.

The attribution maps reveal localized regions of high predictive importance that correspond to The attribution maps reveal that the strongest predictive signal is concentrated around the phosphorylated residue at position 2 of the peptide (Fig. 4B,C). This position corresponds to the primary anchor site in the binding motif of HLA-B*40, where negatively charged residues such as glutamic acid or aspartic acid are typically preferred. The model assigns high attribution to the phosphate group and surrounding structural features of the phosphorylated serine, indicating that the classifier recognizes this modification as a chemically compatible substitute for the canonical anchor residues. In contrast, neighboring positions in the peptide exhibit substantially lower attribution scores, suggesting that the prediction is driven primarily by structural features associated with the anchor position rather than global peptide properties. These observations support the hypothesis that the chemically informed representation enables the model to capture physicochemical similarities between phosphorylated residues and canonical amino acids that fulfill the same structural role in peptide-MHC binding.

Together, these results demonstrate that the chemically informed representation allows the model to recognize functionally equivalent structural features across canonical and modified residues.

### Effects of dataset composition and class imbalance

Interpretation of model performance must take into account the characteristics of the underlying dataset. The peptide-MHC dataset used in this study exhibited substantial class imbalance, with non-binding peptides greatly outnumbering binders. On average, approximately 15% of peptides associated with a given allele were labeled as binders. The distribution of binders and non-binders across the dataset is illustrated in Fig. 5A,B, highlighting the skew toward negative examples. In addition, allele coverage was uneven, with some HLA alleles represented by thousands of peptides while others had comparatively few training examples (Fig. 5C). These factors likely contributed to variability in model performance across alleles and may have limited the ability of the autoencoder to learn stable structural representations for sparsely represented alleles.

**Figure 5.**
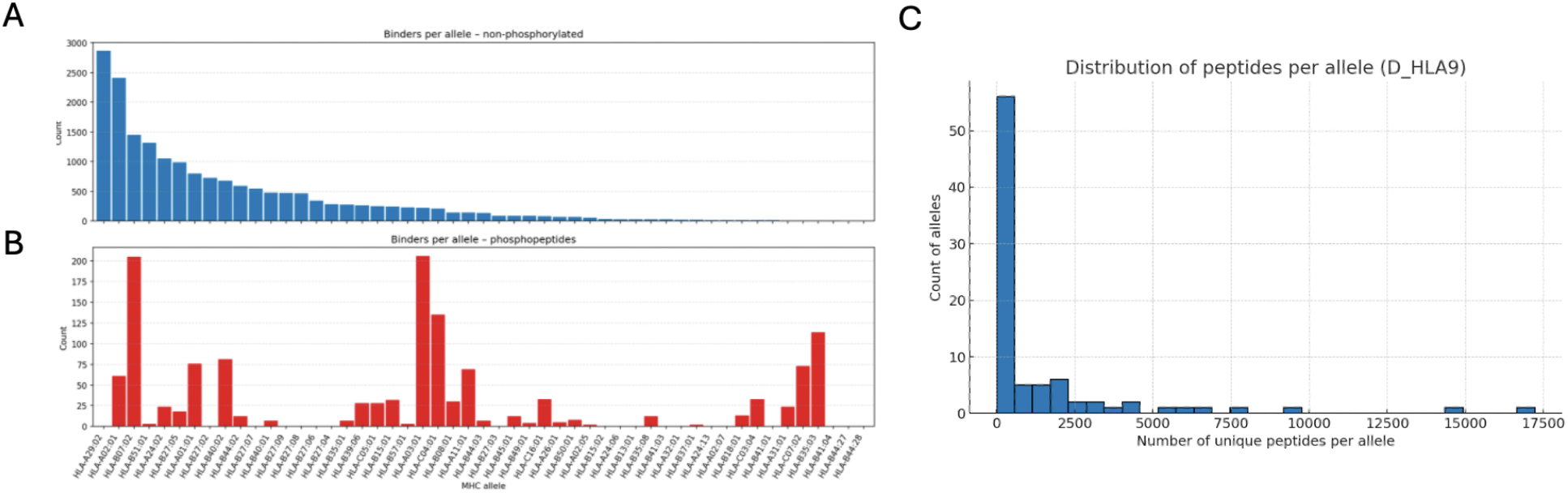
Distribution of peptide binders and dataset composition across HLA alleles. **A**, Number of experimentally observed non-phosphorylated peptide binders per HLA allele. The distribution shows substantial variability in the number of peptides available for different alleles, with a small number of alleles contributing a large fraction of the training data. **B**, Number of phosphorylated peptide binders per HLA allele. In contrast to the non-phosphorylated dataset, phosphopeptide binders are comparatively rare and unevenly distributed across alleles. **C**, Histogram showing the distribution of the total number of unique peptides assigned to each HLA allele in the dataset (D_HLA9). Most alleles are represented by relatively small numbers of peptides, whereas a few alleles contain large peptide collections. This uneven distribution highlights the class imbalance and variability in allele coverage within the dataset used for model training and evaluation.

### Representation behavior on phosphorylated peptides

Because the proposed representation explicitly encodes molecular structure, it naturally accommodates chemically modified residues such as phosphorylated amino acids. Reconstruction experiments demonstrated that when phosphorylated peptides were included in the training data, the autoencoder was able to accurately reproduce the phosphate group within reconstructed peptide images (Fig. 6). However, when phosphorylated residues were rare or absent in the training set, reconstruction quality deteriorated and reconstructed features became more diffuse across the peptide structure (Fig. 6).

**Figure 6.**
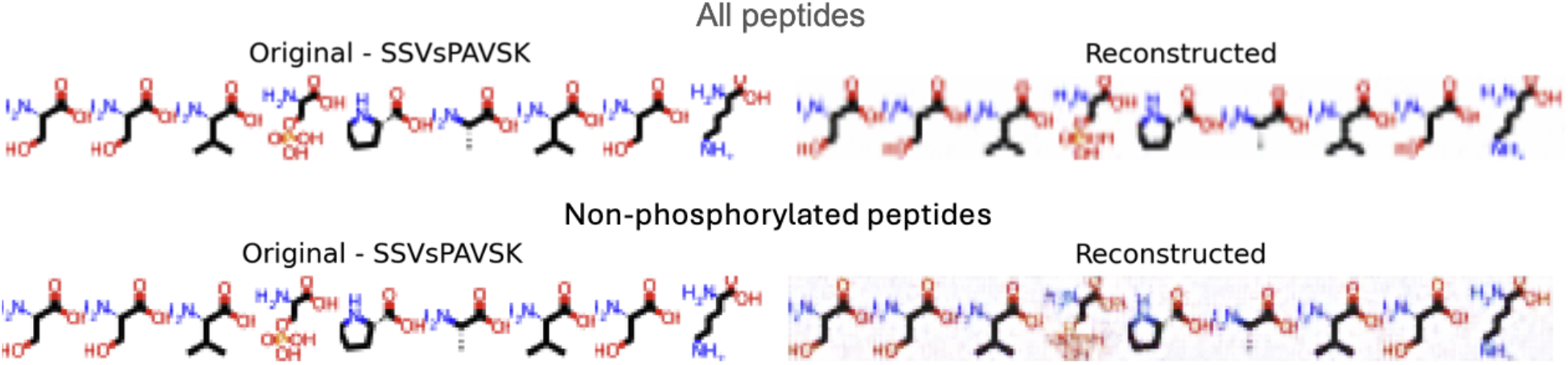
Reconstruction of peptide depictions of a phosphorylated peptide. Mosaic depictions of the peptide SSVsPAVSK and their corresponding reconstructions produced by the convolutional autoencoder. Here the lower case s in position 4 (P4), represents the phosphorylated amino acids. For each peptide, the original peptide depiction (left) is shown alongside the reconstructed image generated by the decoder (right). **Top row (All peptides):** Autoencoder trained on all peptides, reconstruction shows that the autoencoder preserves the overall peptide backbone and side-chain structures in the reconstructed image. **Bottom row (Non-phosphorylated peptides):** Autoencoder trained on non-phosphorylated peptides, the reconstruction shows consistent structural recovery of the phosphate group in P4. These examples illustrate how the learned latent representations capture chemically relevant structural information that can be used in downstream predictive tasks.

Together these observations suggest that while the representation itself can encode chemically modified residues, reliable modeling of such features depends on the availability of sufficient training examples. Importantly, the structural representation provides a flexible framework for incorporating modified residues without introducing additional symbolic tokens into the sequence alphabet.

## Discussion

In this study we explored whether chemically informed representations of amino acids could support machine learning models for peptide analysis. Instead of representing peptides as sequences of symbolic residue identities, we encoded amino acids as two-dimensional depictions of their molecular structures and assembled them into peptide mosaics. A convolutional autoencoder was then used to learn compact latent embeddings from these structural representations. When evaluated in the task of predicting peptide binding to MHC class I molecules, the learned embeddings supported meaningful predictive performance and enabled visual attribution analyses linking model predictions to specific regions of the molecular depictions.

Although models trained on conventional one-hot sequence encodings generally achieved higher predictive performance, the image-based representations retained competitive predictive signal across multiple HLA alleles. This outcome is not unexpected. Peptide-MHC binding is strongly influenced by position-specific anchor residues within the peptide sequence, and one-hot encodings explicitly preserve residue identity at each position. In contrast, the structural representation used here emphasizes physicochemical similarity between residues rather than exact symbolic identity. Given the nature of these two embeddings we expect the autoencoder representation to need more data to achieve a similar performance. However, the chemically informed representation has shown to capture structural and physicochemical properties of residues even when the peptide modification, phosphorylation was not included in the training dataset. This method is the first of its kind and paves the way for an unprecedented approach to learn beyond the 20 amino acid letter code.

Representing amino acids through molecular depictions enables direct interpretation of model behavior using visual attribution methods. Because the input representation corresponds to recognizable chemical structures, attribution maps can be projected onto peptide depictions to identify structural regions that influence predictions. In this study, saliency analyses revealed localized regions of importance corresponding to specific amino acid side chains and positions within the peptide sequence. These observations suggest that the model learns associations between structural features of residues and peptide-MHC binding outcomes. Such interpretability is difficult to achieve with conventional symbolic sequence encodings, where attribution methods highlight sequence positions but not the chemical features responsible for model predictions.

Dataset characteristics also played an important role in model performance. The peptide-MHC dataset used in this study exhibited substantial class imbalance, with non-binding peptides greatly outnumbering binders. In addition, allele coverage varied widely, with some HLA alleles represented by thousands of peptides while others contained relatively few training examples. These factors likely contributed to variability in predictive performance across alleles and may have limited the ability of the autoencoder to learn stable structural representations in sparsely represented datasets. Future studies using larger and more balanced datasets may therefore improve the stability and predictive capacity of chemically informed peptide representations.

A key motivation for this work was the challenge of representing PTMs in sequence-based machine learning models. Conventional protein representations rely on a fixed alphabet of canonical amino acids and therefore require ad hoc extensions to incorporate modified residues. In contrast, the representation proposed here encodes residues through their molecular structure, allowing chemically modified amino acids to be represented directly without introducing new symbolic tokens. Reconstruction experiments demonstrated that phosphorylated residues could be captured within the learned representation when sufficient training examples were available. However, the scarcity of phosphopeptides in the dataset limited the reliability with which such features could be modeled. This highlights an important limitation of current PTM datasets and suggests that improved experimental coverage of modified peptides will be essential for developing robust predictive models of PTM-dependent interactions.

More broadly, chemically informed representations may provide a pathway toward machine learning models that operate on the structural properties of amino acids rather than their symbolic identities. By learning physicochemical relationships between residues, such representations could potentially generalize across structurally related amino acids and chemical modifications. This capability may be particularly valuable in contexts where proteins contain non-canonical residues, chemical modifications, or synthetic peptide designs that fall outside the traditional amino acid alphabet.

Several avenues for future work follow from this study. First, alternative neural architectures such as graph neural networks or attention-based models could be explored to more directly capture relationships between atoms within amino acid structures. Second, pretraining representation models on large collections of peptide depictions could improve the quality of learned embeddings before applying them to specific prediction tasks. Third, richer visual representations, including higher-resolution depictions or encodings that preserve additional structural detail, may enable the model to capture subtler chemical features of amino acids and modified residues. Fourth, integrating sequence-based and chemically informed representations may allow models to combine the strengths of motif-based recognition with structural feature learning. Finally, extending the representation to include additional types of post-translational modifications or non-canonical amino acids may enable broader applications in immunology, protein engineering, and therapeutic peptide design.

Taken together, our results demonstrate that peptides can be represented using chemically explicit structural depictions and that compact embeddings learned from these representations support predictive modeling of peptide-MHC interactions. Although conventional sequence encodings remain highly effective for tasks driven by position-specific motifs, chemically informed representations offer complementary advantages by capturing physicochemical relationships between residues and enabling interpretable analysis of learned features.

Importantly, because these representations encode the underlying chemical structure of amino acids, they provide a framework for incorporating chemically modified residues such as post-translational modifications without expanding the symbolic amino acid alphabet. These findings suggest that integrating structural representations with sequence-based models may provide a promising direction for developing machine learning approaches capable of modeling the broader chemical diversity of proteins. Such chemically informed representations may ultimately enable machine learning models to reason about protein function across the broader chemical space of modified and synthetic amino acids.

## References

(1) 10.1016/0022-2836(70)90057-4

(2) 10.1016/0022-2836(81)90087-5

(3) 10.1016/S0022-2836(05)80360-2

(4) 10.1073/pnas.89.22.10915

(5) Dayhoff, M. O., Schwartz, R. M. & Orcutt, B. C. A model of evolutionary change in proteins. In Atlas of Protein Sequence and Structure, Vol. 5 (Suppl. 3), 345–352 (1978).

(6) 10.1038/s41586-021-03819-2

(7) 10.1038/nbt.3300

(8) 10.1007/s00894-001-0058-5

(9) 10.1002/bip.20296

(10) 10.1073/pnas.2016239118

(11) 10.1126/science.ade2574

(12) 10.1002/pro.2449

(13) 10.1002/prot.22786

(14) 10.1002/anie.200501023

(15) 10.1038/srep00090

(16) 10.1038/nrm1939

(17) 10.1038/nsmb.1842

(18) 10.1038/ncb0502-e127

(19) 10.1146/annurev.bi.61.070192.003553

(20) 10.1093/glycob/cww086

(21) 10.1126/science.1175371

(22) 10.1126/science.ads0018

(23) 10.1016/j.sbi.2025.102997

(24) 10.1093/bioinformatics/btu598

(25) 10.1016/j.immuno.2021.100005

(26) 10.1093/nar/gku1267

(27) 10.1016/j.jaut.2025.103443

(28) 10.64898/2025.12.12.693959

(29) 10.1172/jci.insight.85633

(30) 10.3390/ijms232415803

(31) 10.1126/science.abg2482

(32) 10.1126/science.adg3925

(33) 10.1007/s00018-022-04126-3

(34) 10.1145/3065386

(35) https://arxiv.org/abs/1409.1556

(36) 10.1109/CVPR.2016.90

(37) 10.1109/CVPR.2015.7298594

(38) https://arxiv.org/abs/1312.6114

(39) 10.1021/ci00057a005

(40) 10.5281/zenodo.10398

(41) 10.1074/mcp.M116.063800

